# Realistic basin shape variation modulates local riverine biodiversity via altered connectivity

**DOI:** 10.64898/2026.04.23.720292

**Authors:** J. M. Calabrese, A. B. García-Andrade, Ismail, E. H. Colombo

## Abstract

Understanding the drivers of biodiversity in the world’s rivers, which are known to be hyperdiverse relative to their coverage area, has been an enduring goal in ecology. While regression-based empirical studies have identified a suite of environmental factors that are correlated with riverine fish biodiversity, these insights are often system-specific and inconsistent across regions. In contrast, a more limited body of studies have suggested that network connectivity of rivers affects fish biodiversity by limiting dispersal, and basin shape may modulate these relationships. The few theoretical papers that have explored basin morphology effects have tended to use extreme network shapes and inconsistent methods, thus limiting general insights. Here, we build on these results to demonstrate that river basin morphology, as measured by log aspect ratio, can alter both network connectivity and biodiversity in simulated, all-else-equal scenarios. First, we quantify variation in log aspect ratio across the world’s 100 largest rivers and use this empirical range of shape variation to guide synthetic experiments. In particular, we use Optimal Channel Networks (OCNs) constrained to basins with log aspect ratios within realistic range to study how shape alters connectivity profiles when node number and basin area are held constant. By coupling OCNs with dendritic neutral models, we demonstrate that variation in aspect ratio and concomitant changes in connectivity lead to substantial changes in simulated biodiversity. Finally, we use Earth Mover’s Distance to establish that basin-shape-induced changes in node-level connectivity distributions are predictive of transformations in node-level distributions of *α* and *β* diversity. Overall, elongated basins such as the Mekong River feature lower species richness (*α*-diversity), higher turnover (*β*-diversity), and less variable distributions of both quantities relative to a square reference basin. Furthermore, approximately one third of the world’s largest rivers are elongated enough to potentially feature statistically-detectable, shape-mediated variation in connectivity and biodiversity.

## 1. Introduction

Understanding the drivers of biodiversity in the world’s rivers, particularly for freshwater fishes, has been an enduring goal in aquatic ecology. Freshwater ecosystems in general, and rivers in particular, are well known for being hyperdiverse considering their water volumes relative to marine environments (Abell et al., 2008), and consequently have attracted considerable evolutionary, biogeographic, and ecological study. Many regression-based empirical studies have identified correlates of fish biodiversity in rivers, with variables related to habitat size or capacity (*e*.*g*., discharge: Hugueny, 1989; Oberdoff et al., 1995; Guégan et al., 1998), energy or ecological opportunity (*e*.*g*., net primary productivity: Guégan et al., 1998; Oberdorff et al., 2011), and history (*e*.*g*., geologic and climatic events: Tedesco et al., 2005; Smith et al., 2010) being especially prominent (Shao et al., 2019). However, the results of such efforts can vary substantially depending on regional focus, scale of analysis, and study methodology. So while there is broad agreement that riverine community assemblages arise from complex interactions among environmental conditions, river history, and geomorphological structure (Heino et al., 2015; Tonkin et al., 2017), the overall relative importance of many prominent explanatory variables continues to be debated. Against this backdrop, connectivity and network structure *per se* have begun to gradually be recognized as important drivers of fish population dynamics and community structure (Fagan, 2002; Shao et al., 2019; Lee et al., 2022a), with increased connectivity generally supporting larger population sizes (Sun et al., 2022), broader distributions (Schick and Lindley, 2007), and higher local species richness (Muneepeerakul et al., 2008).

Parallel developments in river geomorphology have deepened our understanding of how drainage basins, and the river networks they contain, are formed. Observations of striking structural regularities across rivers (Horton, 1945; Rinaldo et al., 1992; Rodríguez-Iturbe et al., 1992; Dodds and Rothman, 2000) that may, at first glance, look different (e.g., Zernitz, 1932; Argialas et al., 1988) have led to the hypothesis that a universal mechanism can explain most of the variation in river networks. In particular, Optimal Channel Network (OCN) theory postulates that the physical process of minimizing the total energy expenditure across a river basin is sufficient to produce structurally realistic river networks (Rinaldo et al., 1992; Rodríguez-Iturbe et al., 1992). Recent developments in implementing OCN models (Carraro et al., 2020) have greatly facilitated the study of biological processes, such as dispersal, that operate on river networks (Carraro and Altermatt, 2022; Lee et al., 2022a).

Network connectivity represents a critical nexus between river structure on the one hand, and outcomes of network-constrained flow processes, such as the dispersal of river-obligate organisms, on the other hand (Shao et al., 2019). Modeling studies can reveal how such flow processes interact with networks because they allow a degree of structural manipulation that is not possible in natural systems. Important advances in understanding network-constrained dispersal processes have been made by adapting neutral biodiversity models to run on networks (Muneepeerakul et al., 2007; Economo and Keitt, 2008; Muneepeerakul et al., 2008; Economo and Keitt, 2010; White and Rashleigh, 2012). The pioneering work of Muneepeerakul et al. (2008) established the real-world relevance of dendritic neutral models by showing they could simultaneously capture multiple empirical biodiversity patterns, while Economo and Keitt (2008) were the first to link variation in network structure, via its effect on network connectivity, to dispersal-mediated biodiversity outcomes. In particular, Economo and Keitt (2008) showed that more chain-like networks exhibit longer path lengths and thus lower connectivity, which in turn leads to lower local species richness when compared to branching, tree-like networks. Economo and Keitt (2010) refined these results by demonstrating how network context mediated local biodiversity via its effects on node-level connectivity. White and Rashleigh (2012) focused exclusively on rivers and framed the problem in terms of variation in basin shape. To bracket the range of possibilities, they studied two extreme cases, which included a “high” chain-like network and a “wide” compact tree-like network. Similar to Economo and Keitt (2008), they found that the chain-like network featured lower biodiversity compared to the compact tree, and they inferred that these differences were driven by corresponding differences in network connectivity (White and Rashleigh, 2012). More recently, Carraro and Ho (2025) employed OCNs constrained to a highly elongated basin coupled with a detailed food-web model to illustrate the impact of basin shape on the trophic structure of model communities.

While these results are suggestive in the context of riverine biodiversity, they have important shortcomings. First, previous studies (Economo and Keitt, 2008; White and Rashleigh, 2012; Carraro and Ho, 2025) focused on structural extremes that do not represent the range of basin shape variation observed in nature. So the extent to which empirically realistic changes in basin shape can affect riverine biodiversity via connectivity variation remains an open question. A second consequence of this focus on structural extremes is that none of these studies examined continuous variation in basin shape, connectivity, or biodiversity outcomes. It is therefore not possible to estimate the critical degree of shape variation at which important deviations in these variables begin. Third, two of the three studies used tree networks as their closest approximation to river networks, which are now known to be inferior to self-organized OCNs when representing the structural features of real rivers (Rinaldo et al., 1992; Carraro and Altermatt, 2022). Fourth and finally, these studies used different models to simulate biodiversity and tracked different metrics of river structure, connectivity, and biodiversity, making direct comparison of results difficult.

Here, we build on previous work to link realistic variation in basin shape to changes in both network connectivity and biodiversity. We quantify basin shape in terms of the log aspect ratio (Shelef, 2018; Yi et al., 2018), *R*, which can be readily estimated from data, and leverage a dataset on the world’s 100 largest rivers to inform our analyses. We use a case study of three rivers to highlight particular *R* values, including the minimum (Mekong), maximum (Chad), and the mean (Parnaíba), while using the full range of observed *R* values to study continuous responses of connectivity and biodiversity to shape variation. For each value of *R*, we generate an ensemble of OCNs to capture stochastic variation in network structure, while holding confounding factors like basin area and network size constant and within empirically typical ranges. We quantify node-level connectivity using closeness centrality, which is a standard network connectivity metric (Marchiori and Latora, 2000) that has been shown to strongly correlate with local species richness (Economo and Keitt, 2008). Finally, we use the formal, network adapted neutral model developed by Economo and Keitt (2008) to simulate biodiversity outcomes and track both the *α* (local species richness) and *β* (turnover in species composition) diversity components. We show that variation in *R* translates directly into variation in node-level connectivity profiles, which then manifests as changes in both the node-level *α* and *β* facets of biodiversity. In particular, we find that elongated basins featuring large, negative *R* values, such as the Mekong River, have node-level connectivity distributions that shift to lower values overall, with the formerly highest connectivity values becoming especially compressed. These cases also feature parallel transformations to node-level *α* diversity distributions becoming more concentrated on lower values, and node-level *β* diversity distributions showing the opposite response. We further demonstrate that almost one third of the range of empirical observed *R* values leads to statistically significant transformations in connectivity and biodiversity distributions in our simulations.

## 2 Methods

### 2.1. Basin shape data

We based our analyses on publicly available, precompiled data on the world’s rivers from the *HydroBASINS* dataset (Linke et al., 2019), and selected an intermediate level of spatial resolution (level 6) to represent individual drainage basins. This resolution choice provides a faithful representation of each basin while avoiding artifacts introduced at both coarser and finer scales. In particular, at very coarse resolutions (*e.g*., level 1), polygons aggregate multiple river systems and obscure individual basin geometry, whereas at very fine resolutions (*e.g*., levels ≥10), basins are fragmented into multiple, small catchments whose shapes are erratic and are dominated by local features. We then trimmed river basins to discard any small, disconnected components. This approach resulted in basins that are described by a single polygon whose shape can be studied quantitatively. Finally, we focus our analyses on the 100 largest processed basins (Fig. 1a) to capture the shape variability present in key basins with clear socio-ecological importance, while avoiding potential distortions in interpretation caused by the inclusion of numerous small, erratic basins. Our dataset therefore includes all major basins, such as the Amazon, Congo, Nile, and Mississippi. In addition, the dataset spans a wide range of climatic, geomorphological, and hydrological conditions, providing a robust basis to explore how basin shape relates to underlying environmental and spatial heterogeneity.

**Figure 1.**
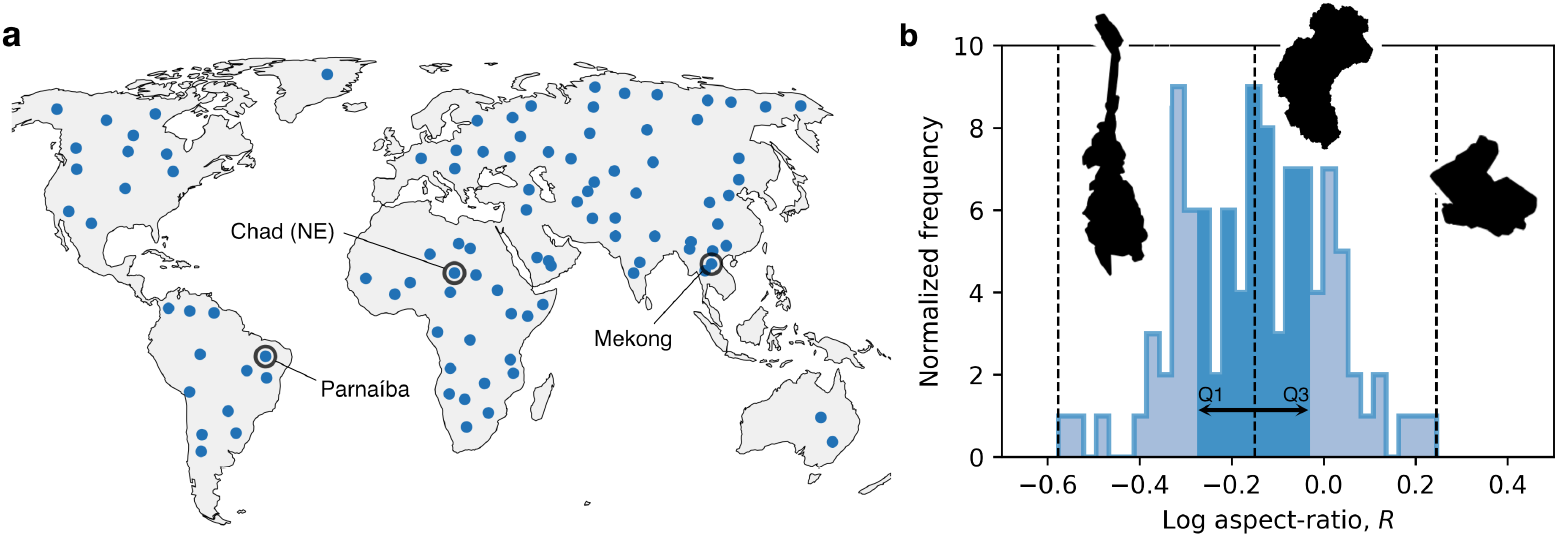
The basin shape dataset for the world’s 100 largest rivers. a) Centroids (blue dots) for the 100 largest basins captured by the *HydroBASINS* dataset at level-6 resolution. Representative case samples are highlighted with black circles: Mekong, Parnaíba, and the North-East portion (outlet-independent) of the Chad basin, corresponding to the minimum, mean, and maximum observed *R* values, respectively. b) Normalized frequency distribution of basin log aspect ratio, *R*. The interval between the first (Q1) and third quartile (Q3) of the distribution is shown in darker blue color, while tails are depicted by a lighter blue tone. Case studies are marked with vertical dashed lines accompanied by their shape (not to scale). In all cases, basin shape is rotated such that its length (*L*_‖_) is vertically aligned and its width oriented horizontally.

### 2.2. Shape characterization

The basin aspect ratio provides a theoretically grounded, first-order characterization of basin shape (Rigon et al., 1996) and is defined by *L*_||_*/L*_⊥_, where *L*_||_ and *L*_⊥_ are the length and the width of the basin, respectively. Given a polygon representing the basin, the length, *L*_||_, can be obtained by calculating the distance from the outlet to the farthest point along the basin perimeter. Basin width, *L*_⊥_, can then be extracted by obtaining the maximum transversal distance relative to *L*_||_. To ensure geometric consistency, these quantities were computed in a projected coordinate reference system chosen to minimize metric distortions. Specifically, we used a locally defined Universal Transverse Mercator (UTM) projection centered on each basin for small to medium basins, and a basin-centered equal-area projection for large basins spanning multiple UTM zones. Finally, taking the square shape as reference, we used the logarithm of the aspect ratio as our descriptor of basin shape,

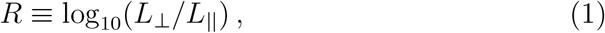

which is conveniently zero for a perfectly square basin, *<* 0 for an elongated basin, and *>* 0 for a wide basin. Importantly, *R* is symmetrical in the sense that, a sign flip (*e.g*., *R* → −*R*) indicates a 90-degree rotation of the basin shape, which is a property not shared by the raw (untransformed) aspect ratio. The histogram of observed *R* values from our 100-river dataset is shown in Figure 1b.

### 2.3. Synthetic basin generation

To focus on how variation in basin aspect ratio alone affects network structure, we generated synthetic basins with user-specified aspect ratio, *r* = *L*_*X*_ */L*_*Y*_, on a rectangular *L*_*X*_-by-*L*_*Y*_ lattice. The side lengths were chosen as integer approximations of 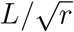and 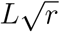 i.e., 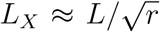 and 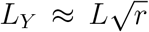, with *L*_*X*_, *L*_*Y*_ ∈ ℤ. For that, we used the recently developed R-package *OCNet* (Carraro et al., 2020), which models the formation of optimal channel networks (OCNs) constrained to the prescribed basin shape. Importantly, OCNs are ideally suited for studying the relationship between basin shape and network connectivity, as they have been shown to reproduce many geometric properties of real rivers (Rinaldo et al., 1992; Rodriguez-Iturbe and Rinaldo, 1997; Carraro and Altermatt, 2022). Networks were considered to be undirected and unweighted, which captures the potential for dispersal along existing waterway connections between different parts of the basin (Muneepeerakul et al., 2008).

To generate OCNs, we set both the total basin area, *A*, and the number of nodes in the network, *N*, to fixed values representative of our dataset. In particular, we set *A* = 10^6^ km^2^, which approximates the mean empirical basin area (≈ 0.75 × 10^6^ km^2^), while the number of nodes was fixed at *N* = 100, taking as reference the mean number of basin subdivisions (sub-basins) at the level-6 resolution (≈ 83 sub-basins). We then configured *OCNet* to generate each basin with a single outlet centered at the bottom and a fixed slope-area scaling exponent (*γ* = 0.5). Additionally, we set the level of noise during OCN formation through the parameters of a cooling schedule and adopted a V-shaped initial condition to enable the network to explore a wide range of structural configurations, ultimately producing realistic river-like patterns (see discussion in Carraro et al. (2020) and Lee et al. (2022b), and details on *Code and data availability*). Finally, we trimmed each synthetic network to remove small, disconnected components, mirroring the preprocessing applied to the empirical basin data.

### 2.4. Network structure and connectivity metric

Structural changes in networks due to shape variation can be quantified by distributions of pairwise distances among nodes (White and Rashleigh, 2012). Such distributions are helpful for visualizing and developing intuition about the broad-scale consequence of network shape variation. However, our main interest lies in quantifying the effect of basin shape changes on node-level biodiversity properties. We therefore require a quantity that aggregates the information contained in the pairwise distance distribution into a node-level descriptor of network structure that can then be directly compared to node-level biodiversity metrics. Node-level connectivity metrics perform this aggregation by using pairwise distances among nodes to quantify the degree to which a focal node is reachable from other points on the network. While many widely-used network connectivity measures exist, we focus here on closeness centrality, *C*, which is a distance-based metric that has been shown to capture the node-level impact of network structure on dispersal processes (see discussion in Economo and Keitt (2010) and Fig. S2). For each OCN-generated network, we calculated the values of closeness centrality, *C*_*i*_, for all nodes, *i*, as,

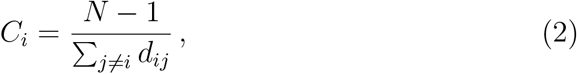

where *d*_*ij*_ is the shortest-path distance between nodes *i* and *j* and *N* is the number of nodes in the network (Marchiori and Latora, 2000).

### 2.5. Dendritic neutral model and biodiversity metrics

We employed a neutral model (Economo and Keitt, 2008) adapted to dendritic networks (hereafter dendritic neutral model; DNM) to quantify the importance of variation in basin-level *R*, and concomitant changes in *C*, for riverine biodiversity processes. Such models have been shown to yield both novel theoretical insights (Economo and Keitt, 2008, 2010) and to be capable of explaining multiple empirical biodiversity patterns in the Mississippi river (Muneepeerakul et al., 2008).

By adapting well-established mathematical tools developed first in population genetics (Kimura, 1991) and later extended to terrestrial ecology (Chave and Leigh, 2002), Economo and Keitt (2008) established a framework to study the dynamics of neutral metacommunities on networks. In particular, they defined an equation for the temporal evolution of the probability, *F*_*ij*_, that two randomly chosen individuals from nodes (sub-basins in our case) *i* and *j* are conspecific,

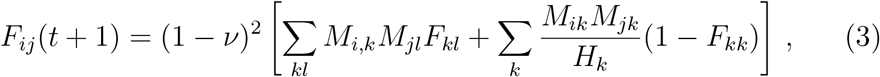

where *v* is the diversification rate, *M* is the migration matrix (flux from *i* to *j*) and *H* is the habitat capacity. As with other neutral metacommunity models (Hubbell, 2011), the model of Economo and Keitt (2008) assumes fixed total metacommunity size with zero-sum dynamics, per-capita equivalence among species in demographic and migration rates, and (infrequent) random speciation.

Equation 3 is solved assuming stationarity: 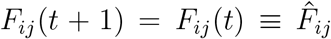. Indices that capture the *α, β*, and *γ* components of biodiversity at node-level can then be defined based on 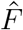 (Economo and Keitt, 2008). Node-level species richness (related to *α* diversity) can be quantified using the Simpson index Simpson (1949), which yields an effective number of species Economo and Keitt (2008),

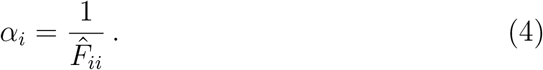

Thus, a low probability that two randomly selected individuals from node *i* are conspecific implies a high effective number of species. We define the node-level *β* component of biodiversity as the average dissimilarity between the community assemblage at a node *i* and those at all other *N* − 1 nodes in the network. Accordingly, for each node *i* we extract

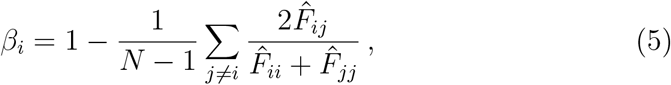

which ranges from 0 (identical assemblages) to 1 (distinct assemblages). Thus, higher *β*_*i*_ indicates greater compositional turnover with the rest of the network.

In its simplest form, the network neutral model of Economo and Keitt (2008) features three user-set parameters: the migration rate *m*, the speciation rate *v*, and the habitat capacity *H*. For our simulations, we set *m* = 0.002, which is an intermediate value that, in combination with a moderate speciation rate of *v* = 10^−5^, allows for connectivity effects to be present but not artificially maximized (see Fig. S1 and Economo and Keitt (2010)). We further assumed, as in Economo and Keitt (2010), that node-level habitat capacity, which governs the number of individuals that can occupy a given node, was homogeneous across all nodes with *H*_*i*_ = 2 × 10^5^. This choice results in a total metapopulation capacity of 2 × 10^7^. Previous results suggest that variation in model parameters can alter the intensity of the observed effects, but does not change them qualitatively (see, e.g., Economo and Keitt (2010) and Fig. S1).

### 2.6. Simulation study design

We choose *R* = 0 as a reference point against which to judge the effects of basin shape variation. Indeed, *R* = 0 falls well within the range of empirical *R* values (−0.6 *< R <* 0.24), studies using OCNs typically default to a square shape (Lee et al., 2022b; Carraro and Altermatt, 2022), and it also represents a geometrically neutral point along the *R* scale that is neither elongated nor compact. For each value of *R*_*k*_, we generated a set of 50 “null” OCNs with *R* = 0, and a set of 50 “alternative” OCNs with *R* = *R*_*k*_. Pooling the 50 “null” OCNs together yields the OCN_0_ dataset, while pooling the 50 “alternative” simulations yields the OCN_*R*_ dataset for the focal value of *R*_*k*_. By applying Eqs. 2, 4 and 5 to each of the OCN datasets, we then extract the corresponding probability distributions of connectivity, *α* diversity, and *β* diversity, respectively, with bin size provided by the Freedman-Diaconis estimator. We use the notation *P*_0_(*Y* ) and *P*_*R*_(*Y* ), to denote the probability distributions of a focal quantity *Y* calculated on the OCN_0_ and OCN_*R*_ datasets, respectively, and where *Y* = *C, α* or *β* depending on the analysis. Comparisons between the probability distributions, *P*_0_ and *P*_*R*_, taken as a function of *R*, for the different quantities of interest allow us to quantify the effects of shape variation on connectivity, *α* diversity, and *β* diversity.

To scan the response of these distributions to changes in basin shape, we generate OCNs with aspect ratio *r*_*k*_ = 0.05, 0.1, …, 2. As a consequence, the log aspect-ratio *R*_*k*_ = log_10_ *r*_*k*_ ranges from approximately −1.3 to 0.3, which encapsulates the empirical range, and extends slightly beyond it in each direction. To streamline the presentation of results and highlight particularly important values of *R*_*k*_, we define a three-river case study by identifying the rivers from our empirical dataset with observed *R* equal to the minimum (Mekong basin, *R* ≈ *R*_5_ = −0.60), mean (Parnaíba basin, *R* ≈ *R*_15_ = −0.15), and maximum (Chad basin, *R* ≈ *R*_33_ = 0.24) values in our sample (for full details see *Code and data availability*). Both the geographic locations, and the positions of the case studies, are shown in Figure 1. This case-study focus is especially important given that many of our results are expressed in terms of probability distributions of focal variables associated with the OCN_0_ or OCN_*R*_ datasets. Furthermore, for most analyses, showing the mean and extreme values of *R* is sufficient to capture the important effects of basin shape on both network structure and biodiversity. Using the whole range of values, *R*_*k*_, we can then perform a systematic investigation to establish threshold values at which basin shape effects become statistically significant (see below), or where resolving a functional relationship between two quantities is necessary.

### 2.7. Distribution transformations

The normalized frequency distributions for connectivity values, *P*_*R*_(*C*), and biodiversity indices, *P*_*R*_(*α*) and *P*_*R*_(*β*) associated with each *R* = *R*_*k*_ serve as metacommunity-level descriptors that can accurately detect distribution transformations across values of *R*. In particular, we quantify (i) how *P*_*R*_(*C*) changes relative to *P*_0_(*C*) as *R* is varied, and (ii) how these connectivity transformations propagate into changes in the *P*_*R*_(*α*) and *P*_*R*_(*β*) biodiversity distributions, relative to *P*_0_(*α*) and *P*_0_(*β*), respectively.

We quantified distributional transformations using the Earth Mover’s Distance (EMD) (Kranstauber et al., 2017), also known as the Wasserstein distance (Kantorovich, 2006), *w*, computed between the baseline (square basin, *R* = 0) and each altered configuration (*R*_*k*_≠0),

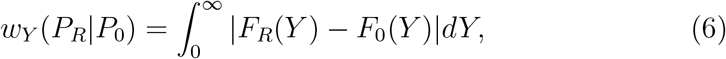

where *F*_0_(*Y* ) and *F*_*R*_(*Y* ) are the cumulative normalized frequency distributions for a focal variable, *Y* = *C, α*, or *β*, and the distributions are evaluated at a given *R* = *R*_*k*_ and the reference case, *R* = 0, as indicated. Selecting the EMD is appropriate because it captures differences between entire distributions rather than just shifts in summary statistics. In particular, *w* is sensitive to changes in shape, spread, and tails, providing a physically interpretable measure of how much “mass” must be redistributed to transform *P*_0_(*Y* ) into *P*_*R*_(*Y* ).

To assess the statistical significance of the observed distance, *w*_*Y*_, we implement a bootstrap procedure to estimate the mean values 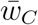, 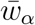 and 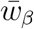, and their associated confidence intervals (Tibshirani and Efron, 1993). More specifically, for each *R*, we repeatedly resample, with replacement, from both the altered distribution *P*_*R*_(*Y* ) and the null distribution *P*_0_(*Y* ), preserving their original sample sizes. For each bootstrap replicate, the distributions are re-estimated on a shared support and the distance *w*_*Y*_ (*P*_*R*_|*P*_0_) is recomputed, yielding a bootstrap distribution of distances from which confidence intervals are obtained via percentile bounds. This facilitates identification of the range of *R* values for which deviations from the null case are statistically significant. To quantify statistical significance relative to the null model, we complement the bootstrap analysis with a Monte Carlo test based on a null distribution of distances. Specifically, we generate this distribution by repeatedly resampling, with replacement, two independent samples from the null distribution *P*_0_(*Y* ), preserving the original sample size, and recomputing the distance *w*_*Y*_ (*P*_0_|*P*_0_) for each replicate. This yields an empirical distribution of distances expected under the null hypothesis of no difference. The p-value is then defined as the proportion of null distances that are greater than or equal to the observed distance *w*_*Y*_ (*P*_*R*_|*P*_0_), with a small finite-sample correction to avoid zero values. This provides a one-sided measure of the probability that an equal or larger deviation could arise under the null. This approach separates the estimation of uncertainty (captured by bootstrap confidence intervals) from the assessment of statistical significance (captured by the null-based p-value).

Finally, to simplify the visualization of the significance and direction of the transformations, we rescale EMD (and associated confidence intervals) to obtain a transformation metric accounting for the intrinsic variability of the null model (i.e., *w*_*Y*_ (*P*_0_|*P*_0_) is not exactly zero due to finite-size effects) signed by the mean displacement,

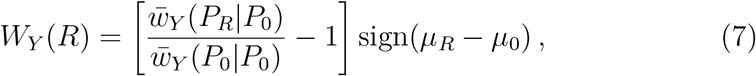

where *µ*_0_ and *µ*_*R*_ are the mean value of *Y* in the null and alternative conditions, respectively. Thereby, with this structure, a deviation with magnitude, |*W*_*Y*_ (*R*)| = 0, which implies 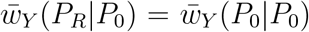, indicates that the altered condition set by *R* generates a distribution of *Y* equivalent to the null condition, while |*W*_*Y*_ (*R*)| *>* 0 indicate the presence of alterations in the distribution profile. Additionally, while the magnitude of *W*_*Y*_ captures the overall transformation of the distribution profile, its sign, based on the displacement of the distribution’s centroid, *µ*_*R*_ − *µ*_0_ (from the null to the altered condition), provides a compact and interpretable description of distribution shift promoted by transformations.

## 3. Results

Our sample of 100 basins spanned a range from highly elongated (Mekong; *R* ≃ −0.6) to moderately compact (Chad; *R* ≃ 0.24), with a mean of *R* ≃ −0.15 (Parnaíba River) and 50% of the cases falling between *R* ≃ −0.27 and *R* ≃ −0.04 (Figure 1). For comparison, the *R* values considered in previous studies of the effects of basin (or network) shape on biodiversity, with one exception noted below, fell outside of the range observed in our sample.

Specifically, the “chain” network considered by Economo and Keitt (2008) has *R* = −∞, the “high tree” studied by White and Rashleigh (2012) has *R* = −1.5 while their “wide tree” network has *R* = 1.0, and the “elongated” case examined by Carraro and Ho (2025) corresponds to *R* ≃ −1.0, while their “compact”, square, case fell inside the empirical range and is equivalent to our null value of *R* = 0.

Examining the distributions of pairwise distances across our three case studies reveals substantial changes for the most negative-*R* observed value, but only modest transformations for the mean and most positive-*R* basins (Figure 2a-c). In particular, for *R* = −0.6 (Mekong case) the distribution has an elongated right tail, with correspondingly larger mean and variance, but a mode that has shifted to the left (Figure 2a). In words, the shortest pairwise distances are unaffected by the change in shape because immediate neighbors remain immediate neighbors. However, path lengths between nodes initially separated by intermediate distances become somewhat longer, while those between nodes that are initially far from each other become substantially longer. Similarly, the Parnaíba case (*R* = −0.15) showed qualitatively similar changes to those in the Mekong case, but with much smaller magnitude (Figure 2b). In contrast, the distribution of pairwise distances in the Chad basin case (*R* = 0.24) showed a somewhat different pattern, with the mode shifting rightwards, but with an overall small magnitude of change compared to the *R* = 0 reference case (Figure 2c). Importantly, transformations in the pairwise distance distributions propagate into *C*, indicating that effectively, closeness can capture the long-range coupling between nodes (beyond first neighbors) that arise due to the spatial neutral dynamics (Figure 2df) (Economo and Keitt, 2010). Transformations in the distributions of *C* were the inverse of those seen in corresponding pairwise distances, but these differences are expected given that the inverse of the sum of all pairwise distance is what enters into the calculation of *C* in equation 2.

**Figure 2.**
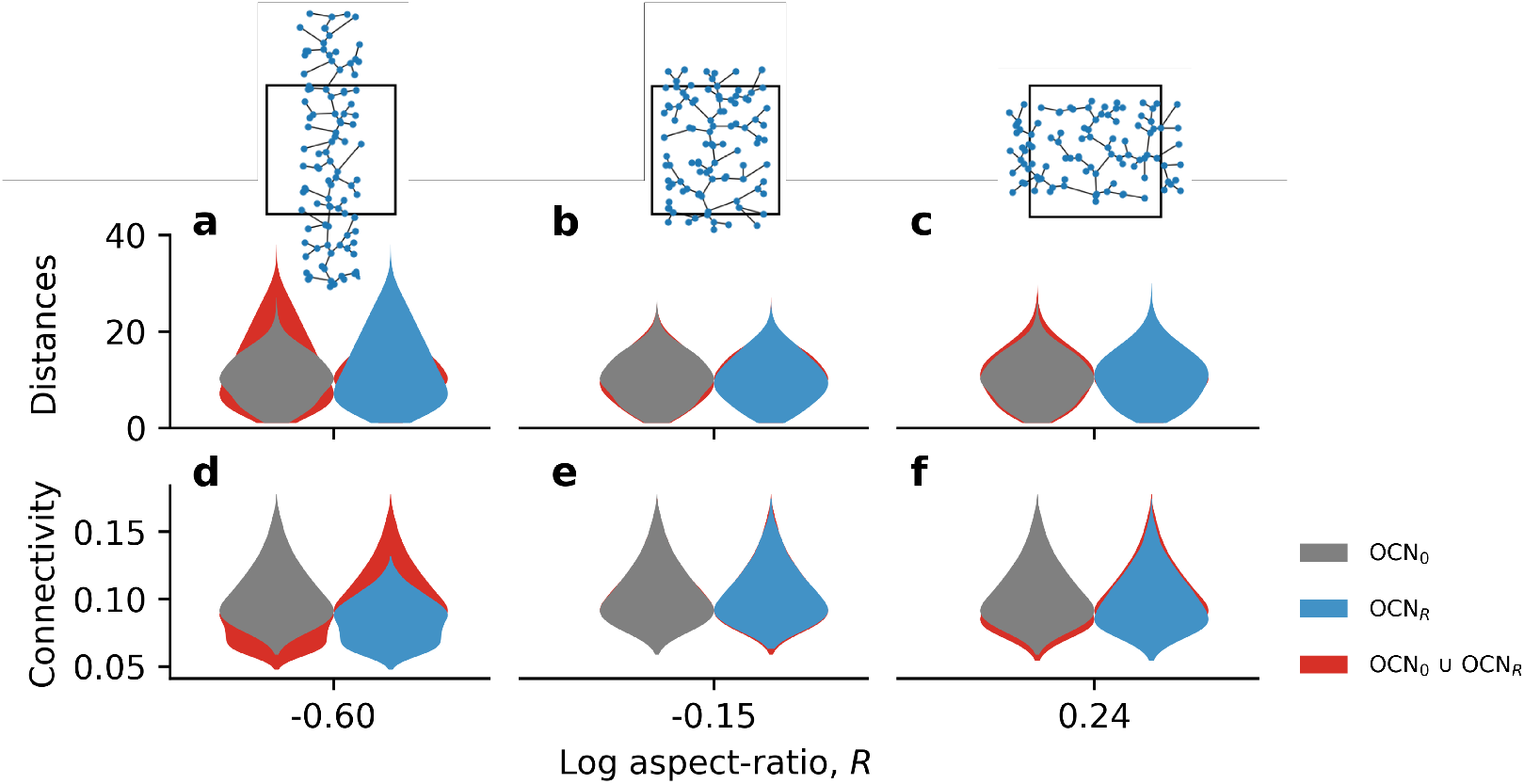
(a-c) Normalized frequency distributions of pairwise distances and (d-f) node-level connectivity for the three case studies. The OCN_*R*_ cases are shown in blue, while the OCN_0_ case is shown in each panel as a black outline for reference. For each representative case, we include an OCN replicate overlapped with an equal-area square for reference. Nodes (sub-basins) are shown in blue dots while water connections are depicted by black solid lines.

More specifically, transformations in the distribution of *C* caused by changes in shape from the *R* = 0 reference case become increasingly pronounced as *R* becomes more negative, while the most extreme positive *R* value observed in our sample had only modest effects on the distribution of connectivity (Figure 3a-c top row). At the most extreme negative *R* value, connectivity values shift downward and become substantially more compressed, with the tail of larger connectivity values being effectively truncated (Figure 3a top row). Crucially, we observe that changes in the distribution of *C* are closely mirrored by changes in the distributions of both *α* and *β* diversity for each representative value of *R* (Figure 3a). Indeed, when measuring distribution transformations between the null (*R* = 0) and alternative (*R*≠0) cases via the EMD, we see that statistically significant transformations in *C* for the *R* = −0.60 (Mekong) case propagate into statistically significant transformations in both the *α* and *β* biodiversity components (Figure 3a). The opposite is also true, with non-significant transformations in the *R* = −0.15 and *R* = 0.24 cases for connectivity being mirrored by non-significant transformations in the distributions of *α* and *β* diversity (Figure 3b, c).

**Figure 3.**
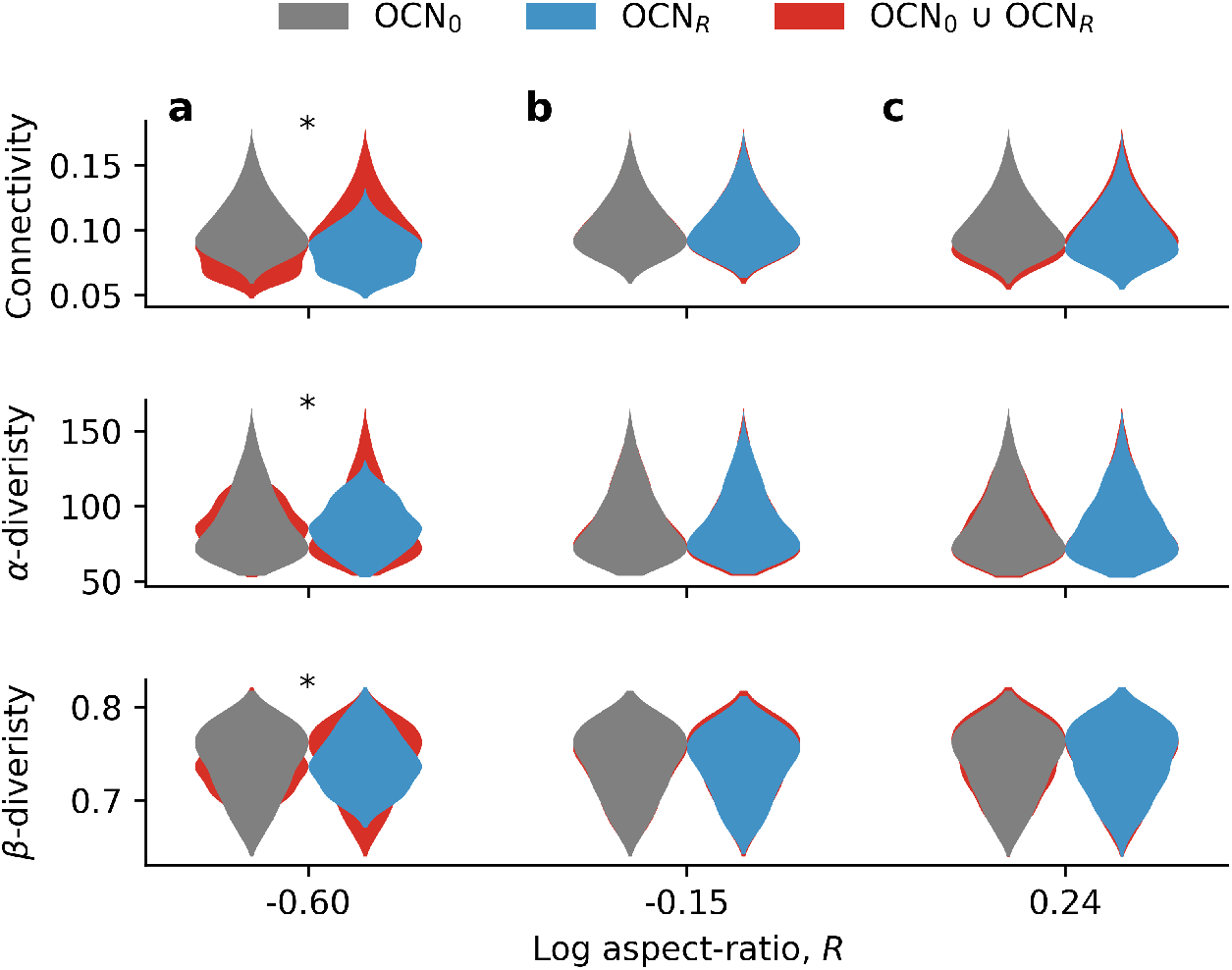
For all three case studies (columns a-c) we show the node-level distributions of connectivity, *α*- and *β*-diversity as violin plots. The OCN_*R*_ based results, *R* ≠ 0, are shown in blue, the OCN_0_ (*R* = 0) results are shown in gray, while the union of both the OCN_0_ and OCN_*R*_ based distributions is shown in red to facilitate comparison of distribution change. Cases where the OCN_*R*_ condition induced statistically significant (*p <* 0.05) distribution transformations are marked with asterisks.

Examining the effect of *R* on shifts in the connectivity distribution in more detail, we see that it increases in magnitude rapidly as *R* becomes more negative (Figure 4a). In particular, the brown shaded region in Figure 4a indicates the *R* value at which the OCN_*R*_ distributions of connectivity, *α* diversity, and *β* diversity all become statistically significantly different from their OCN_0_ counterparts. This region includes 32% of the 100 basins, meaning that almost a third of the world’s basins could potentially exhibit basin-shape signatures in connectivity and, in turn, biodiversity. We further find that connectivity transformations, as quantified by the EMD, are highly predictive of both *α* (Fig. 4b) and *β* (Fig. 4c) biodiversity transformations, with most values clustering near the ±1:1 line in both cases. As the EMD quantifies changes in shape across the entire distributions, it is sensitive not just to induced shifts in central tendency, but also to changes in the skewness and in the asymptotic behavior of the distribution (Kantorovich, 2006), this confirms that these distributions all shift in lock step in response to variation in *R*.

**Figure 4.**
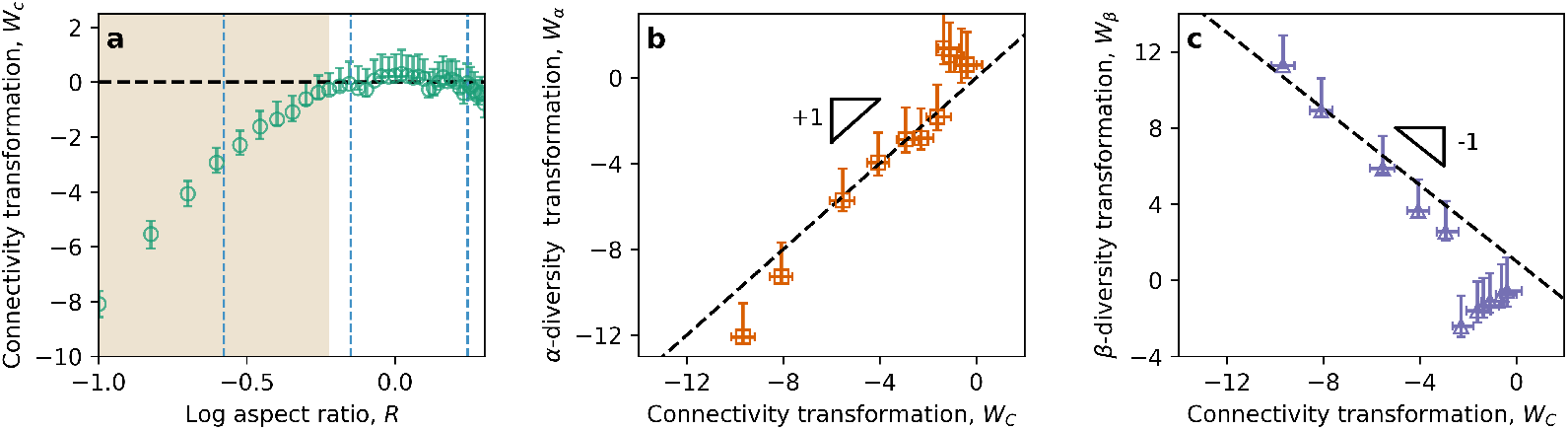
a) Connectivity transformation metric, *W*_*C*_, as a function of *R*. b) *α*- and c) *β*-diversity transformations, *W*_*α*_ and *W*_*β*_, respectively, as a function of connectivity transformation. Light brown region in panel a) indicate the range of *R* where transformations for all three variables are statistically significant. In panels b and c, we include reference lines with slopes +1 and −1 for reference (black dashed lines).

Finally, we show that both *α* and *β* diversity are highly correlated with *C*, with the former biodiversity metric being positively related and the latter being negatively correlated (Figure 5). While the relationship between each biodiversity statistic and connectivity weakens slightly and becomes somewhat compressed at the most extreme negative *R* value (Figure 5a-c), the overall relationship is not qualitatively affected. The stability of the relationship between these variables is a direct consequence of the tightly coupled way that the distributions of *C, α* diversity, and *β* diversity all shift together in response to variation in *R*.

**Figure 5.**
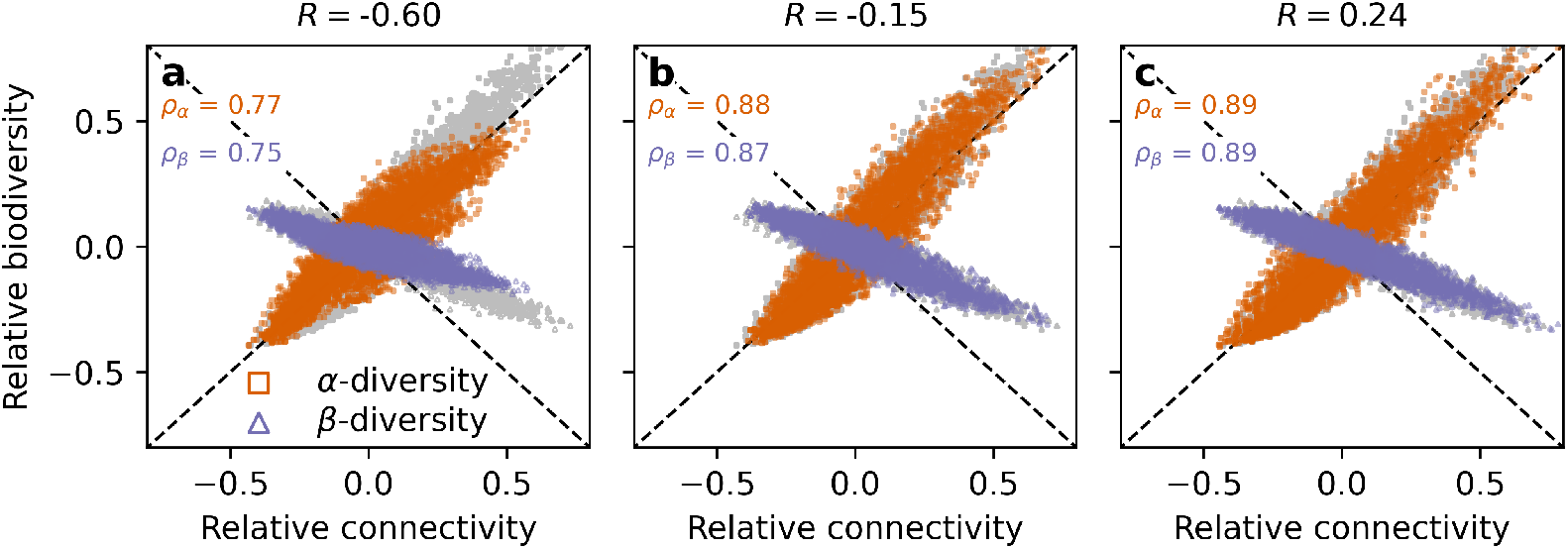
Both biodiversity metrics are strongly and approximately linearly correlated with connectivity across the range of *R* captured by the case studies. The OCN_*R*_ associated values are shown in purple for *α* diversity and orange for *β* diversity. The corresponding OCN_0_ values are shown in gray in the background for each case to provide an additional visual reference for judging how variation in *R* affects the correlation between connectivity and biodiversity. To facilitate visualization of the biodiversity response to connectivity variation we show the value of each variable relative to its mean (and then shifted to zero), e.g. for *C* we plot 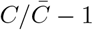, where 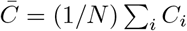. The origin of each plot (0, 0) then corresponds to the nodes that have the average connectivity and average biodiversity index values. Dashed positive (for *α* diversity) and negative (for *β* diversity) 1:1 lines are shown for reference.

## 4. Discussion

We have shown that realistic variation in *R* can profoundly alter the distributions of both *α* and *β* diversity across a basin via its effects on network connectivity. In particular, we have demonstrated that very elongated basins like the Mekong River (*R* = −0.6) tend to have overall lower *α* diversity and higher *β* diversity compared to an otherwise-identical square reference case (*R* = 0). In both cases, these changes in mean biodiversity are accompanied by corresponding contractions in the distributions of biodiversity metrics, implying that biodiversity metrics tend to be less variable across nodes in more elongated basins. While the strength of these effects accelerates with increasingly negative *R* values, transformations in the within-basin distributions of both biodiversity metrics were already statistically significant at *R* = −0.26 (Rio Grande basin), which is within the inner-quartile range of observed *R* values. The location of this significance threshold implies that almost a third of the world’s basins could potentially show statistically detectable, shape-induced biodiversity change. Though our simulations show that the effects of *R* on both connectivity and biodiversity are roughly symmetrical for a given absolute value of *R*, real river basins are strongly skewed towards negative log aspect ratios. Elongation is indirectly implied (Rigon et al., 1996) by the empirical observation of Hack’s law (Hack, 1957), where the longest stream length scales superlinearly with area, *L* = *A*^*h*^, with *h >* 1*/*2. This indicates that length increases faster with area when compared to an isotropic scaling (*h* = 1*/*2) (Rigon et al., 1996). Elongation therefore emerges from the same physical and organizational principles governing river network structure, and has direct consequences for the distribution of biodiversity within the basin. While other studies have inferred that shape affects biodiversity via connectivity (Economo and Keitt, 2010; White and Rashleigh, 2012; Carraro and Ho, 2025), here, we more directly establish this causal link and assess whether and to what extent it becomes significant across the range of shapes observed in natural basins. In particular, we created all-else-equal scenarios where we only varied *R* within the same class of network, explicitly quantified the distributions of node-level connectivity and corresponding distributions of node-level biodiversity metrics, and then used the EMD to show that transformations in the distribution of connectivity strongly predict transformations in the distributions of both biodiversity components. It is important to recognize here that, within our modeling framework, causality between connectivity and biodiversity can only flow in one direction. To see this, consider that our metric of connectivity, Closeness Centrality, is dependent only on network structure and is completely unaffected by changes in biodiversity in the system. The converse, however, is not true. Instead, changes in connectivity clearly affect the distribution in biodiversity, and the EMD quantifies this dependency. This result highlights the tight coupling between connectivity change and biodiversity change and shows that the within-basin distributions of all three quantities move in lock step in response to variation in *R*.

Despite its strengths, our approach still has important limitations that must be recognized. While dendritic neutral models are elegant, simple, and have been shown to be real-world relevant (Muneepeerakul et al., 2008), their empirical support is currently restricted to one well-known example, and whether or not such results will generalize across basins remains to be seen. Second, while neutral models emphasize fundamental demographic processes including dispersal, they are, by design, silent with respect to issues such as trophic structure, functional traits, and niche differentiation. When more detailed insights into community structure are required, such as understanding foodweb composition (Carraro and Ho, 2025) or quantifying distributions of functional traits (Su et al., 2023), more elaborate models will need to be employed. A further limitation is that current implementations of OCNs only allow variation in morphology in a very limited way, namely as variation in the rectangular bounding area of the basin. While we have accommodated this limitation by considering our aspect ratio metric relative to the square, baseline case for OCNs, real basins are unquestionably more complex in shape and morphology. For example, higher-order aspects of basin morphology, such as the smoothness of basin boundaries, may also engender variation in both connectivity and biodiversity, but unfortunately can’t be studied in a systematic way with currently available OCN-focused methods.

We have established that changes in basin shape alone, within the real-world range of basin morphology, cause changes in the distribution of node-level connectivity within the resulting river networks. We have further shown that these connectivity changes propagate into highly predictable changes in the node-level distributions of both *α* and *β* diversity. These results suggest some interesting topics for future study. First, while we have held network size constant here, many structural properties of river networks are known to scale allometrically with system size. Importantly, these scaling laws are not linear in basin area, as observed in the case of Hack’s law (*e.g*., length of longest stream: Hack, 1957) and Flint’s law (*e.g*., basin slope: Flint, 1974). While Colombo et al. (2025) made a first exploration of connectivity scaling in dendritic networks, our results showing the effects of basin shape on connectivity, and subsequently, on biodiversity, suggest that network size alone may not sufficiently explain variation in network connectivity across basins varying in size, shape, and other factors. Finally, though the effects of connectivity on riverine biodiversity are beginning to be recognized, global-scale studies of the connectivity-biodiversity relationship in rivers are still lacking. Such studies could lend much-needed perspective on the importance of connectivity in biodiversity determination relative to well-established environmental factors.

## Acknowledgments

This work was partially funded by the Center of Advanced Systems Understanding (CASUS), which is financed by Germany’s Federal Ministry of Research, Technology and Space (BMFTR) and by the Saxon Ministry for Science, Culture and Tourism (SMWK) with tax funds on the basis of the budget approved by the Saxon State Parliament.

## 6. Code and data availability

The code and supporting data used in this study are publicly available in the GitHub repository *basinShape* (Colombo, 2026). The repository includes all scripts required to reproduce the main computational results and figures presented in this work. The datasets provided are a compiled and processed version of *HydroBASINS* Level 6 hydrographic data, which serves as the underlying raw source (Linke et al., 2019). Access to *HydroBASINS* Level 6 data is available through its official distribution platform.

