## Supplementary Figures for "Realistic basin shape variation modulates local riverine biodiversity via altered connectivity"

### Supplementary material for:
### Realistic basin shape variation modulates local riverine biodiversity via altered connectivity

#### I. SUPPLEMENTARY FIGURES

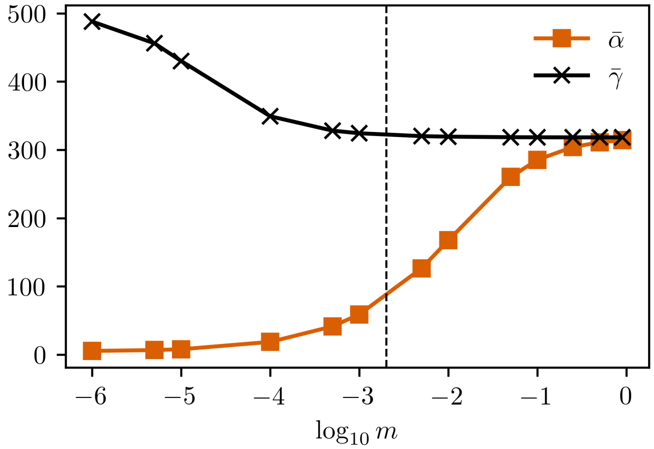


Figure S1. Average node-level species richness, $\bar{\alpha} = (1/N)\sum_i \alpha_i$ and metacommunity total species richness, $\bar{\gamma} = (1/N^2)\sum_{(i,j)1/F_{ij}}$ (see Economo and Keitt [1]) for increasing migration rates. For these results we used an elongated basin (Mekong case, $R = -0.6$). The intermediary migration rate we used in the main text is highlighted with vertical dashed line (for further details see *Materials and methods* section in the main document).

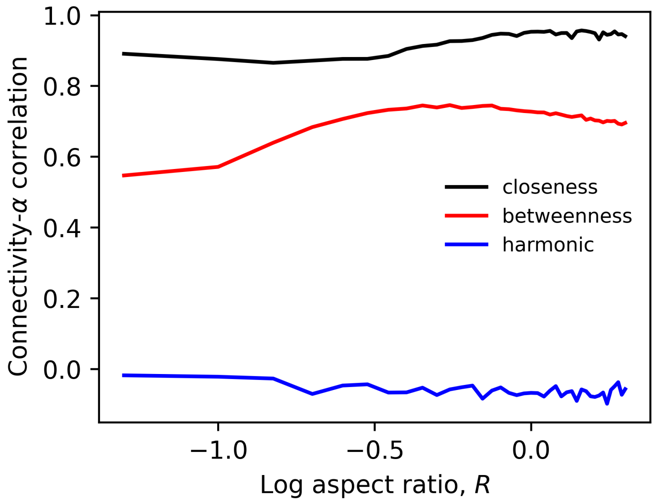


Figure S2. Pearson correlation coefficient between connectivity and $\alpha$-diversity metric as a function of log aspect ratio, $R$ (details are described in the main text).

[1] E. P. Economo and T. H. Keitt, Species diversity in neutral metacommunities: a network approach, Ecology Letters **11**, 52 (2008).
